# Nuclear plasticity regulates macrophage phagocytic capacity *in vivo*

**DOI:** 10.64898/2026.01.22.700862

**Authors:** Clelia Amato, Rosalind Heron, Will Wood

## Abstract

Macrophages fine-tune their appetite to fulfil their clearance role, but how they achieve this is incompletely understood. The nucleus can sense and respond to extracellular physical challenges, but whether it can also detect and react to intracellular mechanical inputs is unclear. Here, we tested the hypothesis that phagosomes exert a strain on the nucleus, and that the consequent nuclear mechanoresponse influences macrophage appetite. Combining genetic manipulation and live *in vivo* imaging, we show that engulfed apoptotic bodies indent the nucleus of *Drosophila* embryonic macrophages, leading to nuclear deformation and causing an increase in Lamin B levels. We demonstrate that these phagosome-induced nuclear molecular and mechanical adaptations are critical for macrophages to sustain uptake – unveiling a role for nuclear plasticity in the regulation of macrophage phagocytic capacity *in vivo*.

## Introduction

Phagocytosis allows the removal of unwanted solid particles, and is critical to maintain and restore homeostasis during development and throughout adulthood. As professional phagocytes, macrophages can internalise numerous targets – sequentially and often simultaneously. Alongside this remarkable appetite, macrophages possess the ability to timely cease uptake. Doing so not only prevents them from reaching the point of lysis^1^, but also safeguards their responsiveness to unprogrammed targets. In fact, overburdened macrophages (often referred to as “exhausted” or “fatigued”) – which are commonly observed in conditions like atherosclerosis^2^ – exhibit reduced motility in response to inflammatory cues^3^ and impaired clearance of bacteria and dead cells at wounds^3,4^. How macrophages regulate their phagocytic capacity has been investigated for more than three decades, but – despite this long-standing interest –plasma membrane availability remains the only confirmed limiting factor^1,5,6^. Simply put, our current understanding is that macrophages halt phagocytosis once their deployable membrane reservoir – which mostly entails recycling endosomes^7,8^ – has been exhausted, meaning that the membrane surrounding internalised cargoes can no longer be replaced at the cell’s edge and extending further phagocytic projections becomes mechanically unfavourable. These studies clearly demonstrate that physical challenges profoundly affect macrophages’ phagocytic behaviour. One, so far largely overlooked, mechanical challenge macrophages face during phagocytosis is intracellular crowding. Following engulfment, the cargo is drawn and accommodated inside the phagocyte’s cytoplasm – where it is likely to collide with pre-existing compartments. These internalisation-driven intracellular mechanical perturbations have been considered inconsequential and largely overlooked; yet the macrophage cytoplasm is equipped with mechanosensitive organelles that may be able to both sense and respond to them. Amongst these is the large, relatively stiff, genome-housing nucleus, which has a well-established role in detecting and reacting to extracellular physical inputs such as spatial confinement^9^, greatly influencing immune cell decision-making^10^. Whether the nucleus senses and responds to intracellular mechanical triggers remains critically understudied^11^ and essentially unexplored in the context of immune cells. Here, for the first time, we looked at phagocytosis as an intracellular force-generating process – testing the hypothesis that internalised material mechanically perturb the nucleus, and that nuclear plasticity is an important regulator of macrophage phagocytic capacity. We did so *in vivo*, exploiting the genetic and live imaging potential of the *Drosophila melanogaster* embryo^12^. We found that internalised apoptotic bodies indent the nucleus of embryonic macrophages, leading the organelle to progressively deform as the number of internalised apoptotic corpses increases during embryogenesis. By genetically removing apoptosis – which results in macrophages reaching the end of embryonic development with no internalised bodies – we demonstrated that phagocytosed apoptotic corpses are not only the direct cause of nuclear deformation but also of adaptive changes in nuclear lamina composition. In fact, we discovered that late-stage corpse-deprived macrophages exhibit a noticeable reduction in Lamin Dm0 (orthologue of human Lamin B) levels, which inevitably impacts the nucleus’ mechanical properties. We established that phagosome-induced molecular adaptations within the nuclear lamina are critical for macrophages to regulate their phagocytic capacity. In fact, preventing internalisation-driven increase in Lamin Dm0 deposition (by knocking down *LamDm0* specifically within macrophages) causes a dramatic reduction in phagocytic load. Collectively, our findings uncover a role for the nucleus in sensing and responding to internalised biomass, ultimately regulating macrophages’ phagocytic capacity *in vivo*.

## Results

### Macrophage nuclear deformation increases alongside phagocytic burden and nuclear indentation during *Drosophila* embryogenesis

To understand whether internalised material mechanically perturb the phagocyte nucleus *in vivo*, we live-imaged *Drosophila* macrophages co-expressing a nuclear (Srp-H2A-3x-mCherry) and a cytosolic (Srp-Gal4.2, UAS-2x-eGFP) marker during early and late embryogenesis. Uptake of apoptotic bodies (which are generated upon physiological turnover of the central nervous system) begins at stage 12 – when macrophages start their migratory journey from the head mesoderm^13^ – and is largely complete by stage 15 – when macrophages have fully dispersed along the embryonic ventral midline (**Figure 1A, Movie S1**). Accordingly, stage 12 macrophages contain fewer bodies (visualised as cytoplasmic voids and henceforth referred to as “vacuoles”) than stage 15 macrophages (**Figure 1B, C**) – with a significantly larger proportion of them containing none (**Figure S1A, Movie S2**). Therefore, the developing embryo is an ideal and physiologically relevant experimental setup to assess the impact of engulfed particles on nuclear dynamics. As shown in **Figure 1D**, stage 15 macrophages appear more prone to dramatic nuclear shape changes than stage 12 macrophages. To quantitate this, we measured the Feret diameter (i.e., the longest distance between any two points along the selection boundary) of individual nuclei, and calculated its standard deviation across 14 minute-long timelapses. This confirmed that the amplitude of nuclear shape change is greater in late-stage than in early-stage macrophages (**Figure 1E, Movie S3**). Interestingly, we found that that macrophages’ nuclear size increases as development proceeds, with stage 15 macrophages possessing a significantly larger nuclear volume (**Figure S1B**). We then asked whether the increase in internalised apoptotic bodies translates into an increase in phagosomes directly contacting the nucleus. To answer this, we scored the number and percentage of indenting vacuoles in early- and late-stage macrophages. We observed that stage 12 macrophages exhibit a significantly lower number of indenting vacuoles than stage 15 macrophages (**Figure 1F**). However, when we normalised this for the total number of internalised debris, we found no difference in the percentage of indenting vacuoles between the two stages (**Figure 1G**) – which suggests that nuclear indentation occurs early during embryogenesis. Further supporting this notion, we found that the vast majority (88.298% ± 4.961, Mean ± S.D., N= 5 embryos, 81 macrophages) of corpse-bearing stage 12 macrophages contain at least one vacuole directly in contact with the nucleus (**Figure S1C**). Overall, our data suggests that internalised apoptotic bodies mechanically perturb the nucleus, and that increasing phagocytic load and nuclear indentation correlate with progressively more pronounced nuclear deformations.

**Figure 1.**
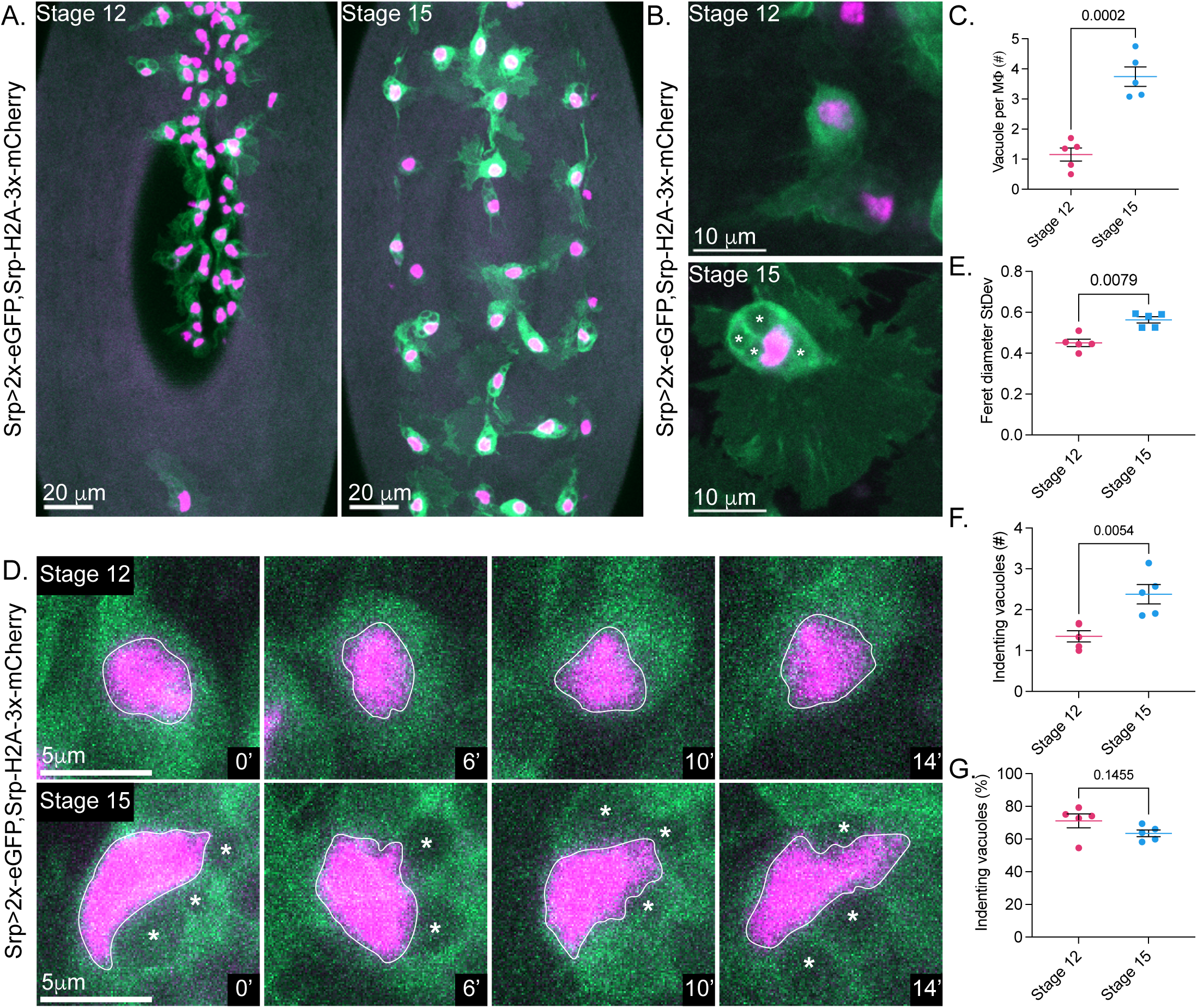
Macrophage nuclear deformation increases alongside phagocytic burden during *Drosophila* embryogenesis. **A.** *In vivo* confocal images of representative stage 12 (left) and 15 (right) embryos expressing a nuclear (Srp-H2A-3x-mCherry, magenta) and a cytosolic (Srp-Gal4.2, UAS-2x-eGFP, green) MΦ marker. **B.** *In vivo* confocal images of a representative stage 12 (top) and 15 (bottom) MΦ expressing a nuclear and a cytosolic marker (as per A.); asterisks: vacuoles. **C.** Quantitative analysis of vacuole number (#) per MΦ. **D.** *In vivo* confocal time-series of representative stage 12 (top) and 15 (bottom) MΦs expressing a nuclear and a cytosolic marker (as per A.); elapsed time (minutes) is indicated at the bottom-right corner of each panel; asterisks: vacuoles. **E.** Quantitative analysis of MΦ nuclear deformation amplitude, expressed as the Feret diameter standard deviation (StDev) across 14 minute-long timelapses. **F.** Quantitative analysis of indenting vacuole number (#) per MΦ. **G.** Quantitative analysis of indenting vacuole percentage (%) per MΦ.

### Phagocytosed apoptotic bodies directly cause macrophage nuclear deformation

We then sought to understand whether the increased phagocytic burden simply correlates with, or is instead the leading cause of, nuclear deformation. To do so, we compared macrophage nuclear shape change in control and *H99* mutant embryos expressing a nuclear (Srp-H2A-3x-mCherry) and a cytosolic (Srp-Gal4.2, UAS-2x-eGFP) macrophage marker. *H99* embryos lack the three master regulators of apoptosis^14^, meaning that embryonic macrophages do not encounter any dead cells to engulf during their migratory journey. As shown in **Figure 2A** and previously reported^15^, macrophages in *H99* embryos undergo normal developmental dispersal, but reach embryonic stage 15 with zero engulfed apoptotic bodies. A visual comparison between equally developed (both stage 15) control and *H99* embryos revealed a striking difference in macrophage nuclear morphology, with phagocytes within *H99* embryos exhibiting a much more regularly shaped nucleus than controls (**Figure 2B**). Live imaging confirmed that macrophages within *H99* embryos maintain their nuclear morphology remarkably unaltered over time, in stark contrast with the rapidly changing control nuclei (**Figure 2C, Movie S4**). Accordingly, the Feret diameter standard deviation of macrophage nuclei within *H99* embryos is significantly lower than that of control nuclei (**Figure 2D**), despite no measurable difference in nuclear volume (**Figure 2E**). Overall, our findings demonstrate that internalised apoptotic bodies are the leading cause of nuclear deformation in embryonic macrophages.

**Figure 2.**
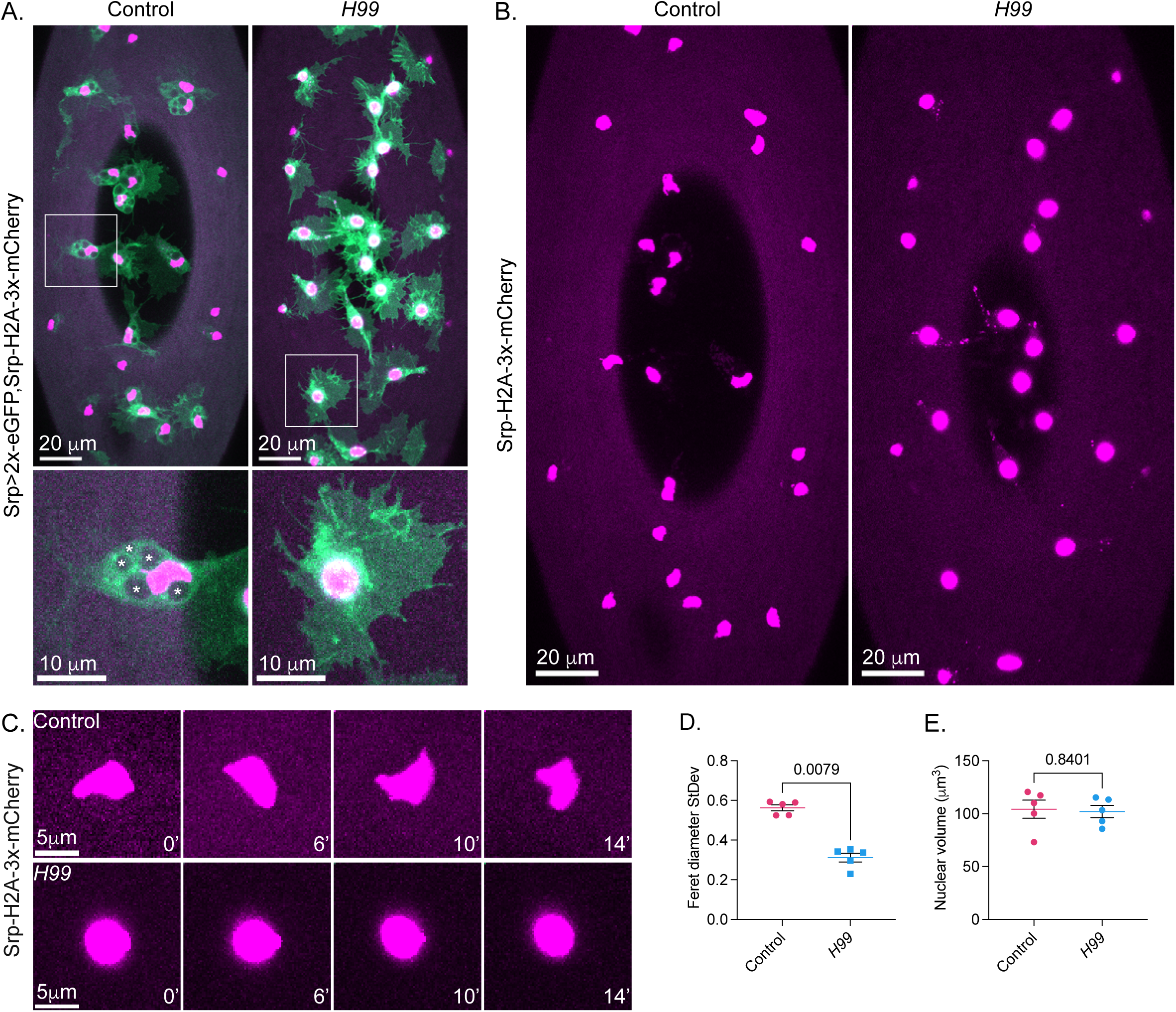
Macrophage nuclear deformation is caused by internalised apoptotic bodies. **A.** *In vivo* confocal images of representative stage 15 control (left) and *H99* (right) embryos expressing a nuclear (Srp-H2A-3x-mCherry, magenta) and a cytosolic (Srp-2x-eGFP, green) MΦ marker; for both conditions, a representative MΦ (square) is shown in greater detail in the inset below. **B.** Single channel (Srp-H2A-3x-mCherry, magenta) confocal images of the same control and *H99* embryos shown in A. **C.** *In vivo* confocal time-series of a representative control (top) and *H99* (bottom) stage 15 MΦ expressing a nuclear marker (as per B.); elapsed time (minutes) is indicated at the bottom-right corner of each panel. **D.** Quantitative analysis of MΦ nuclear deformation amplitude, expressed as the Feret diameter standard deviation (StDev) across 14 minute-long timelapses. **E.** Quantitative analysis of MΦ nuclear volume.

### Phagocytosed apoptotic bodies alter nuclear lamina composition

Nuclear morphology is largely dependent on the nuclear lamina, a meshwork of intermediate filament proteins (i.e., Lamins) lying underneath the inner leaflet of the nuclear envelope. We therefore sought to understand whether the more regular nucleus observed in macrophages within *H99* embryos present obvious differences in nuclear lamina composition in comparison to controls. Uniquely among invertebrates, and more similarly to higher organisms, the *Drosophila* genome harbours two Lamin genes – *LamDm0* and *LamC* – which encode the orthologue of human Lamin B and Lamin A/C respectively^16^. The generally accepted view is that Lamin B contributes to nuclear elasticity^17^, Lamin A/C determines nuclear stiffness^18^, and the stoichiometric ratio between the two dictates nuclear rigidity^19^. However, our current understanding of the relative contribution of Lamins to immune cell nuclear mechanics is significantly less than that of other cell types^20^. Since *LamDm0* is constitutively expressed – unlike *LamC*, which becomes transcriptionally active at later stages of embryonic development ^16^ – we reasoned that changes in Lamin Dm0 levels were more likely to account for any measurable difference in nuclear lamina composition between controls and macrophages within *H99* embryos. To gain insights into the physiological levels of Lamin Dm0, and compare them to that of macrophages within *H99* embryos, we performed immunofluorescence experiments using an anti-Lamin Dm0 antibody. To unequivocally distinguish macrophages from other Lamin Dm0-positive cells – such as epidermal cells, which highly express *LamDm0* (**Figure 3A**) and are in close z-proximity to our phagocytes of interest (**Movie S5**) – we fixed and stained embryos expressing a nuclear (Srp-H2A-3x-mCherry) and a cytosolic (Srp-Gal4.2, UAS-2x-eGFP) macrophage marker. As shown in **Figure 3B**, late-stage embryonic macrophages express detectable levels of Lamin Dm0 within their nuclear lamina. When we compared equally developed (both stage 15) control and *H99* embryos, we found that macrophages within apoptotic-deficient embryos possess lower levels of Lamin Dm0 (**Figure 3C**), which we confirmed by quantitative analysis (**Figure 3D**). Overall, our data demonstrates that internalised apoptotic bodies profoundly change nuclear lamina composition in embryonic macrophages.

**Figure 3.**
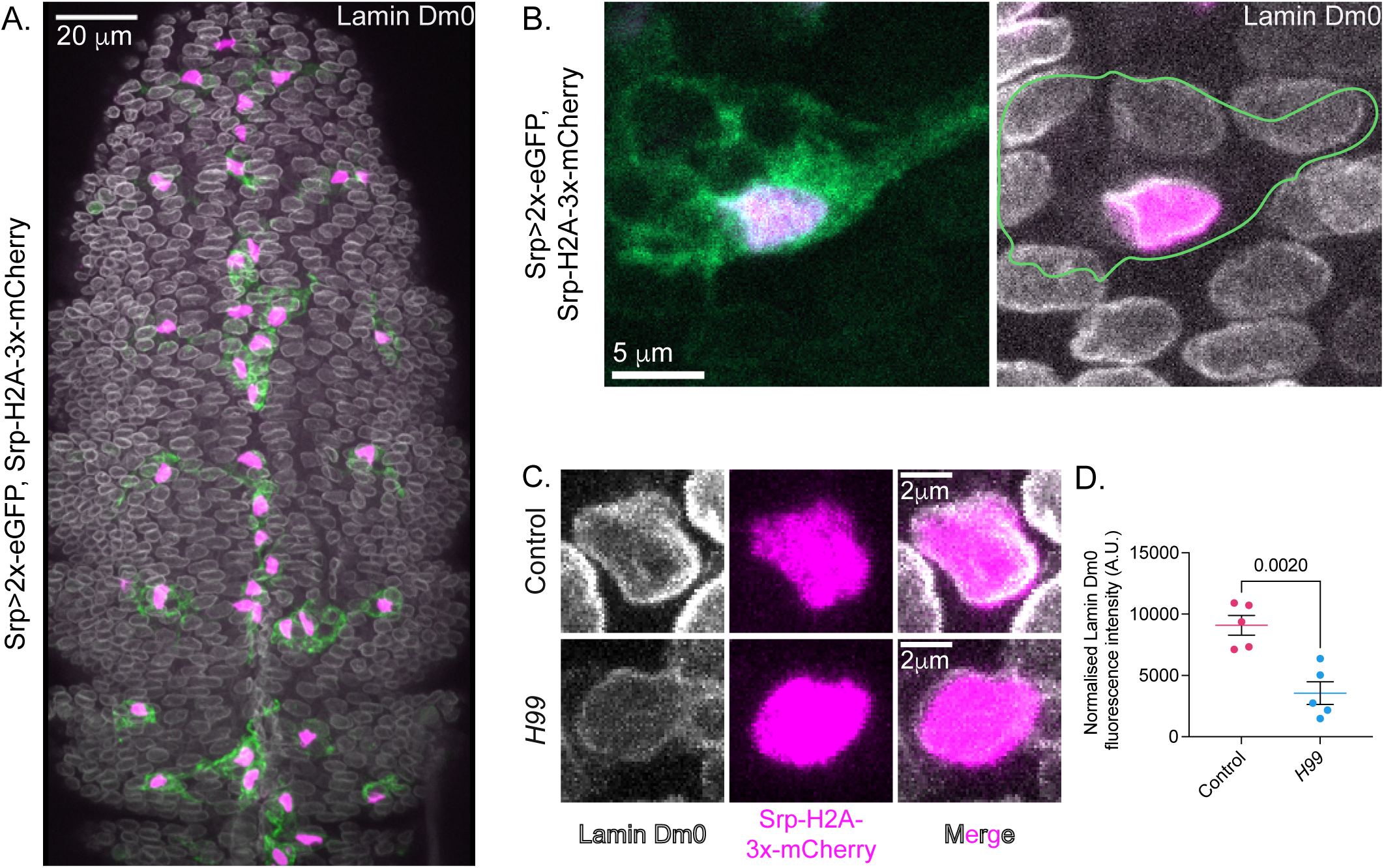
Internalised apoptotic bodies cause an increase in Lamin Dm0 levels. **A.** Confocal image of a representative control stage 15 embryo expressing a nuclear (Srp-H2A-3x-mCherry, magenta) and a cytosolic (Srp-2x-eGFP, green) MΦ marker fixed and stained with an anti-Lamin Dm0 antibody (grey). **B.** Confocal image of a representative stage 15 control MΦ showing a nuclear and a cytosolic marker (as per A., left), and a nuclear marker (as per A., green channel not shown) alongside the anti-Lamin Dm0 staining (grey, right). **C.** Confocal images of a representative stage 15 control MΦ (top) and *H99* mutant MΦ (bottom) expressing a nuclear (Srp-H2A-3x-mCherry, magenta) and a cytosolic (Srp-2x-eGFP, not shown) marker (as per A.) fixed and stained with an anti-Lamin Dm0 antibody (grey). **D.** Quantitative analysis of Lamin Dm0 fluorescence intensity (normalised for background fluorescence) in MΦs within control and *H99* embryos.

### Preventing internalisation-driven molecular adaptation of the nuclear lamina causes a reduction in macrophage phagocytic load

Finding that late-stage vacuole-deficient macrophages (i.e., within *H99* embryos) possess lower levels of Lamin Dm0 than equally developed controls suggests that – under normal circumstances – phagocytosis leads to an increase in Lamin Dm0 deposition. We hypothesised that adaptive changes in nuclear lamina and, consequently, in nuclear mechanical properties may be critical for macrophages to withstand vacuole indentation as this becomes more prominent during embryogenesis (Figure 1F-G). To test this hypothesis, we knocked-down *LamDm0* specifically in macrophages using two independent, Srp-Gal4.2-driven, RNAi lines – henceforth referred to as *LamDm0* RNAi #1 (which has recently been used to down-regulate *LamDm0* in pupal macrophages^21^) and *LamDm0* RNAi #2. As shown in **Figure 4A**, loss of Lamin Dm0 does not compromise macrophage developmental dispersal but does affect their morphology, with *LamDm0* knock-down phagocytes exhibiting less noticeable vacuoles than controls. Quantification of vacuole number and size revealed that, despite no significant difference in the overall number of vacuoles per macrophage (**Figure 4B**), *LamDm0* knock-down phagocytes possess smaller vacuoles than controls (**Figure 4C**). Strikingly, unlike vacuoles in control macrophages – which remain visible in maximum intensity projections – the smaller vacuoles within *LamDm0* knock-down macrophages, visible within a single optical plane, often become hidden in projected images (**Figure 4D**). We next asked whether loss of Lamin Dm0 negatively affects direct interactions between phagosomes and nucleus. To answer this question, we quantified nuclear indentation in Lamin Dm0-depleted macrophages. As shown in **Figure 4E**, loss of Lamin Dm0 significantly reduces direct contacts between the vacuoles and the nucleus, with the smaller vacuoles observed in *LamDm0* knock-down phagocytes frequently observed in peripheral regions of the cell body (Figure 4D). Lastly, given that nuclear indentation correlates with nuclear deformation (Figure 1), we sought to understand whether loss of Lamin Dm0 affects nuclear shape change. To do so, we compared the Feret diameter standard deviation of control and *LamDm0* knock-down macrophages. As shown in **Figure 4F** and quantified in **Figure 4G**, the nucleus of Lamin Dm0-depleted macrophages is as deformable as control nuclei despite major changes in lamina composition. Intriguingly, in agreement with recent work^21^, we also found that *LamDm0* knock-down macrophages possess a smaller nucleus than controls (**Figure 4H**). Overall, our data indicates that preventing phagosome-induced molecular adaptation within the nuclear lamina dramatically reduces macrophages’ phagocytic capacity.

**Figure 4.**
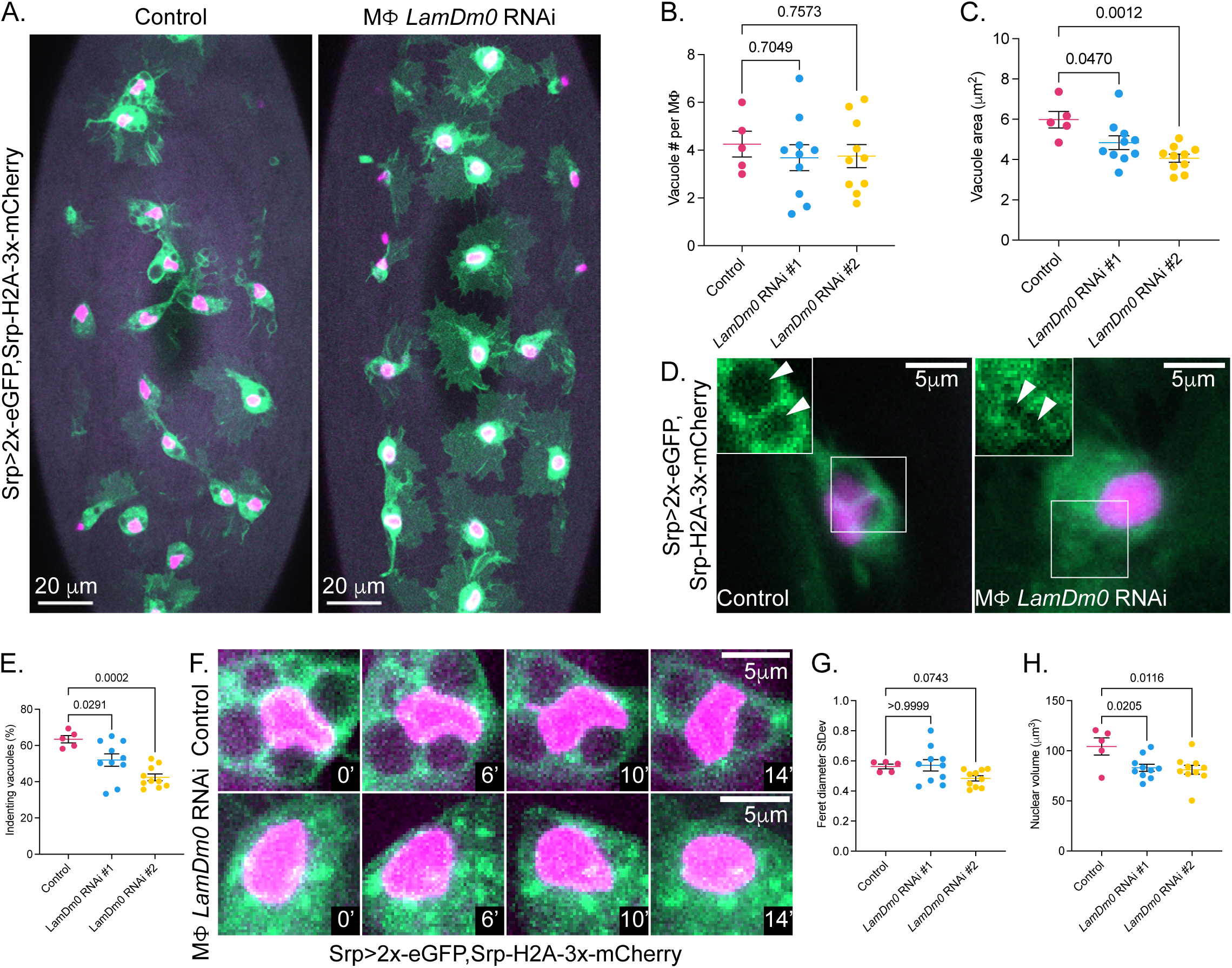
Preventing internalisation-driven nuclear adaptation causes a reduction in macrophage phagocytic load. **A.** Representative *in vivo* confocal images of control (left) and MΦ *LamDm0* RNAi (right) stage 15 embryos expressing a nuclear (Srp-H2A-3x-mCherry, magenta) and a cytosolic (Srp-2x-eGFP, green) MΦ marker. **B.** Quantitative analysis of vacuole number (#) per MΦ in control and *LamDm0* RNAi (#1 and #2) MΦs. **C.** Quantitative analysis of vacuole area per MΦ in control and *LamDm0* RNAi (#1 and #2) MΦs. **D.** *In vivo* confocal images (maximum intensity projections) of a representative control (left) and MΦ *LamDm0* RNAi (right) MΦ expressing a nuclear and a cytosolic marker (as per A.); vacuoles at their maximal size (square) are shown as individual focal planes within the insets (arrowheads). **E.** Quantitative analysis of the percentage (%) of indenting vacuoles in control and *LamDm0* RNAi (#1 and #2) MΦs. **F.** *In vivo* confocal time-point of representative control (top) and MΦ *LamDm0* RNAi (bottom) MΦs expressing a nuclear and a cytosolic marker (as per A.); elapsed time is indicated at the bottom-right corner of each panel. **G.** Quantitative analysis of nuclear deformation amplitude, expressed as the Feret diameter standard deviation (StDev) across 14 minute-long timelapses, in control and *LamDm0* RNAi (#1 and #2) MΦs. **H.** Quantitative analysis of nuclear volume in control and *LamDm0* RNAi (#1 and #2) MΦs.

## Discussion

As macrophages patrol the organism, they constantly assess their surroundings – discerning “eat me” from “don’t eat me” signals to unequivocally spot targets for phagocytosis. Whilst some of these targets are a staple diet for macrophages (e.g., those derived from events of programmed cell death), others are less commonly encountered (e.g., pathogens and necrotic debris at sites of tissue damage). To ensure that unexpected debris are effectively removed, macrophages limit their phagocytic capacity to carefully avoid steady-state overburdening. By doing so, they remain motile (to reach targets located outwith their immediate proximity, if necessary) and maintain sufficient accommodating space inside their cytoplasm. Although a thorough characterisation of macrophage intracellular crowding has yet to be undertaken, we anticipate that the conclusions reached by an elegant study – which explored the spatial distribution and interaction network of essentially omnipresent organelles (e.g., nucleus, endoplasmic reticulum, Golgi apparatus, mitochondria)^22^ – can be extended to our phagocytes of interest. Despite an incredibly low “free space” inside their body, macrophages can make room for many – often very large – particles. We started from the simple but largely overlooked notion that phagocytosis exacerbates cytoplasmic crowding, unavoidably leading to intracellular collisions, to test two key hypotheses: 1) Macrophages possess (at least) one mechanism to infer their cytoplasmic compaction; 2) they use it to control their appetite. We envisioned the nucleus as a plausible sensor for cytoplasmic crowding for several different reasons. First, it is a large organelle – occupying approximately one third (30.8% ± 8.4%; Mean ± S.D.; N: 5 embryos, 102 macrophages) of the cell surface of *Drosophila* embryonic macrophages, which we have used here as an *in vivo* experimental system. Second, it is a relatively stiff compartment, up to 10 times stiffer than the surrounding cytoplasm ^23^ – and, as such, rate-limiting in different cellular processes, most notably spatially confined migration ^24,25^. Third, it can sense and respond to extracellular mechanical inputs (with physical confinement again being the most extensively studied) and impact immune cell behaviour^10,21^. Exploiting the elegant genetics and powerful *in vivo* live-imaging potential of the *Drosophila* embryo, we first established that internalised apoptotic corpses mechanically perturb the macrophage nucleus. Specifically, we found that – immediately after being phagocytosed – apoptotic bodies indent and progressively deform the nucleus as embryogenesis proceeds. Accordingly, we also found that late-stage corpse-deprived macrophages within apoptosis-deficient embryos (i.e., *H99* mutants) exhibit a substantially undeformed nucleus. Given that these cells migrate along the narrow channel formed between the epidermis and the central nervous system without undergoing notable nuclear shape changes, our results argue that – *in vivo* – phagocytosis exerts an even more potent mechanical strain on the nucleus than the widely studied spatially confined migration. Furthermore, we discovered that early uptake/indentation trigger an adaptive molecular change to the macrophage nuclear lamina – similar to the confinement-induced adaptation recently described in pupal macrophages^21^. Specifically, we measured an increase in Lamin Dm0 (orthologue of human Lamin B) deposition in the nuclear lamina of vacuole-rich macrophages. Being able to trigger such molecular (and, consequentially, mechanical) adaptation turned out to be essential for macrophages to successfully sustain further rounds of internalisation. Indeed, macrophage-specific knock-down of *LamDm0* led to a dramatic reduction in their phagocytic capacity – resulting in drastically smaller phagocytic vacuoles. Importantly, we also found that loss of Lamin Dm0 reduces nuclear indentation, which – we propose – occurs in the attempt to relieve compaction upon a nucleus with compromised molecular and mechanical plasticity. Phagosome perinuclear localisation has previously been reported, and proposed to be required for lysosomal fusion and cargo degradation^26^. Our study goes one step further, demonstrating that phagosome intracellular positioning does not only affect its own fate but also that of targets yet to be internalised. Taken together, our results lead to a beautifully simple model whereby, *in vivo*, early phagocytic events lead to nuclear indentation/deformation, causing molecular and mechanical adaptations to the nuclear lamina that allows macrophages to withstand further phagocytic events. In lay terms, we think about the earliest internalised apoptotic bodies as anticipating raindrops, prompting the nucleus to change into a more appropriate – waterproof – gear to effectively handle a storm. Our work represents a major advance in our understanding of how macrophages regulate their phagocytic capacity. As macrophages become increasingly more promising therapeutic “tools and targets”^27^, further insights into their engulfing potential is essential to harness their therapeutic potential fully. More broadly, our work provides an entirely novel perspective to look at how cells – arguably not only immune types – interpret, and are influenced by, intracellular mechanical cues.

## Acknowledgements

We are immensely grateful to Luke Tweedy, PhD for generating the macro required for nuclear volume measurements. We would like to acknowledge Helen Falconer (University of Edinburgh) for her critical work, and the Bloomington Stock Center (University of Indiana, NIH) for providing *Drosophila* lines. We are thankful to members of the Wood lab and the Edinburgh Cell Death Collective (ECDC) for helpful discussions.

## Author contributions

Conceptualization: CA, WW

Methodology: CA, WW

Investigation: CA, RH

Formal analysis: CA

Visualization: CA, WW

Funding acquisition: CA, RH, WW

Writing – original draft: CA, WW

Writing – review & editing: CA, WW

## Data and materials availability

*Drosophila* strains generated or in this study are available upon request.

## Supplementary Materials

### Material and methods

#### Drosophila husbandry

All *Drosophila* strains used for this work (listed in Table S1) were kept at 18°C, room temperature (21°C) or 25°C (with 50-60% relative humidity in a 12:12h light/dark cycle) on standard cornmeal-agar food. Prior to embryo collection, adult flies were put in laying cages with apple juice agar plates spread with yeast paste.

### Embryo preparation for live imaging and fixation/immunostaining

Embryos were collected from overnight laying cages and transferred to cell strainers (Falcon), dechorionated in bleach (Jangro) for 90 seconds and washed six times in distilled water.

For live imaging, dechorionated embryos were manually selected: the desired genotype was identified by the presence of fluorescent markers and the correct developmental stage by gut morphology. Selected embryos were mounted ventral side up on a glass slide coated with double sided sticky tape, and a No 1.5 coverslip (SLS) was added to the slide’s left and right margins. Embryos were embedded in VOLTALEF oil (VWR), covered with a No 1.0 coverslip (SLS), sealed and left to dry for 10 minutes prior to imaging.

For fixation/immunostaining, dechorionated embryos were transferred in a glass vial containing a 1:1 solution of 4% Formaldehyde and heptane, shaken for 30 seconds and incubated for 30 minutes on a rotating wheel at room temperature. The fixative was then removed and 100% methanol added to the glass vials, followed by a vigorous 30 second shake. Devitellinised embryos were transferred to a new glass vial and rinsed three times with 100% methanol. Embryos were washed three times with PBS- 0.3% TritonX (Tx)- 0.5% BSA (PBS-Tx-BSA) and incubated overnight at 4° C with primary anti-Lamin Dm0 antibody (ADL67.10, Developmental Studies Hybridoma Bank; 5 μg/ml). Stained embryos were washed three times in PBS-Tx-BSA and blocked using 0.2% Horse Serum in PBS-Tx-BSA for 30 minutes at room temperature. Embryos were then incubated with a secondary 405 goat anti-mouse antibody (A31553, Thermo Fisher Scientific; 1:100 in PBS-Tx-BSA) for 1 hour at room temperature, washed three times in PBS-Tx-BSA and embedded in VECTASHIELD medium (H-1000, Vector Laboratories). In preparation for imaging, embryos were selected on a glass slide, and desired embryos transferred on a new glass slide coated with double sided sticky tape and containing a No 1.5 coverslip (SLS) on the margins. Embryos were embedded in VOLTALEF oil, covered with a No 1.0 coverslip (SLS), sealed and left to dry for 10 minutes prior to imaging.

### Live and fixed/stained embryo imaging

Live and fixed/stained embryos were imaged using a Zeiss LSM980 confocal microscope equipped with a 40x/1.3 oil immersion objective. For both live and fixed embryos, GFP and mCherry were excited using a 488 and 561 nm laser respectively; for stained embryos, endogenous Lamin Dm0 was imaged using a 405 laser. To quantitate nuclear deformability, time-lapses were acquired at a 0.58 μm interval over an average depth of 28.39 μm (for stage 12 embryos) or of 17.18 μm (for stage 15 embryos) every 60 seconds. To quantitate Lamin Dm0 levels, three equally sized areas were imaged per embryo at a 0.45 μm interval over an average depth of 30.05 μm.

### Image processing & analysis

Maximum intensity projections were generated using the Zeiss Zen Blue software, exported as .czi files and imported to Fiji^28^. Otherwise, unprojected images were exported as .czi files and transformed into maximum or average intensity projections in Fiji. The Fiji 2.14.0/1.54f version was used to then process the images as needed (e.g., cropping and linear brightness/contrast adjustment).

### Phagocytic vacuoles number and area

To quantitate phagocytic vacuoles in stage 12 vs stage 15 control MΦs, the number of cytoplasmic voids per MΦs was counted on maximum intensity projections and averaged per embryo using Microsoft Excel. To quantitate phagocytic vacuoles in control and *LamDm0* RNAi MΦs, the number of cytoplasmic voids per MΦs was counted on individual focal planes and averaged per embryo using Microsoft Excel. The area of control and *LamDm0* RNAi MΦ phagocytic vacuoles was measured in Fiji using the freehand selection tool, manually outlining each cytoplasmic void on the focal plane corresponding to its maximal extension. Obtained values were then exported to Microsoft Excel to calculate the average phagocytic vacuoles area per MΦ and then averaged for each embryo.

### Nuclear deformation

To measure the amplitude of nuclear shape change, embryos were imaged for 14 minutes at a regular 60-second interval. Individual nuclei were outlined manually using the freehand selection tool or the Wand (tracing) tool (Tolerance: 100) in Fiji at all time-points on maximum intensity projections. The Feret diameter was measured in Fiji, obtained values exported to excel to calculate the Feret diameter standard deviation for each macrophage and then averaged for each embryo.

### Nuclear indentation

To measure the number of indenting vacuoles, cytoplasmic voids directly in contact with the nucleus were scored on individual focal planes for each MΦ. The average number of indenting vacuoles was calculated using Microsoft Excel, and obtained values were averaged per embryo. To measure the percentage of indenting vacuoles, the number of phagocytic vacuoles directly in contact with the nucleus was divided by the total number of cytoplasmic voids within each MΦ, and obtained values were then averaged per embryo.

### Nuclear volume

To measure nuclear volume, we used the Fiji 3D Objects Counter function^29^. This required the following pre-processing steps, which generated a binary stack. 1) We applied a 3D median filter over neighbouring pixels to increase the robustness of boundary detection in the next step. 2) We used the Otsu method to threshold the stacks, as this maximises the variance between the foreground and background, and is robust to differences in brightness from stack to stack. We found it to reliably differentiate between the nuclei and the background after median filtering. 3) Because some nuclei are in contact, they can appear to be a single particle. To correct for this, we watershed the image.

### LamDm0 endogenous levels

To quantitate the endogenous levels of lamin Dm0, three equally sized (1024 x 1024 pixels) areas corresponding to the anterior (A), central (B) and posterior (C) ends were imaged per embryo. To measure Lamin Dm0 levels within each MΦ, A,B or C images were cropped (135 x 135 pixels) to include only one MΦ nucleus, and a maximum intensity projection was generated including only the focal planes where each MΦ nucleus was visible. Each nucleus was manually outlined using the freehand selection tool in Fiji (5-pixel width), and the mean intensity fluorescence for the appropriate channel measured. The obtained values were then normalised by subtracting the background fluorescence, calculated as the average fluorescence (relative to same channel) across a three straight 2.52 pixels long lines.

### Statistical analysis

To ensure that the appropriate statistical tests were performed, all datasets underwent Shapiro-Wilk normality tests. Unpaired t-tests and Mann-Whitney tests were then performed on normally distributed and non-normally distributed data respectively. Datasets with more than two groups were compared using a one-way ANOVA test or Kruskal-Wallis test, as recommended by the GraphPad Prism (Version 10.4.0) software. All graphs show mean ± SEM. Statistical details can be found in Table S2.

## Supplementary Figures

**Figure S1.**
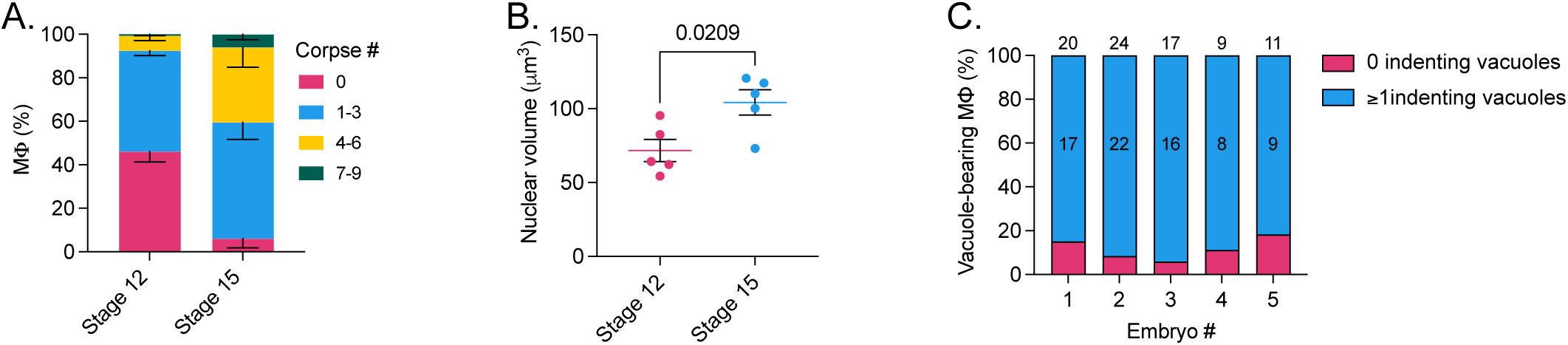
Vacuolation, nuclear size and phagosome indentation in embryonic macrophages. **A.** Quantitative analysis of the percentage (%) of MΦs with 0, 1-3, 4-6 and 7-9 internalised apoptotic bodies in stage 12 and stage 15 control embryos. **B.** Quantitative analysis of nuclear volume in stage 12 and stage 15 control MΦs**. C.** Percentage (%) of corpse-bearing stage 12 control MΦs possessing at least 1 vacuole indenting the nucleus: Embryo 1: 17 out of 20, Embryo 2: 22 out of 24, Embryo 3: 16 out of 17, Embryo 4: 8 out of 9, Embryo 5: 9 out of 11.

## Supplementary tables

**Table S1.**
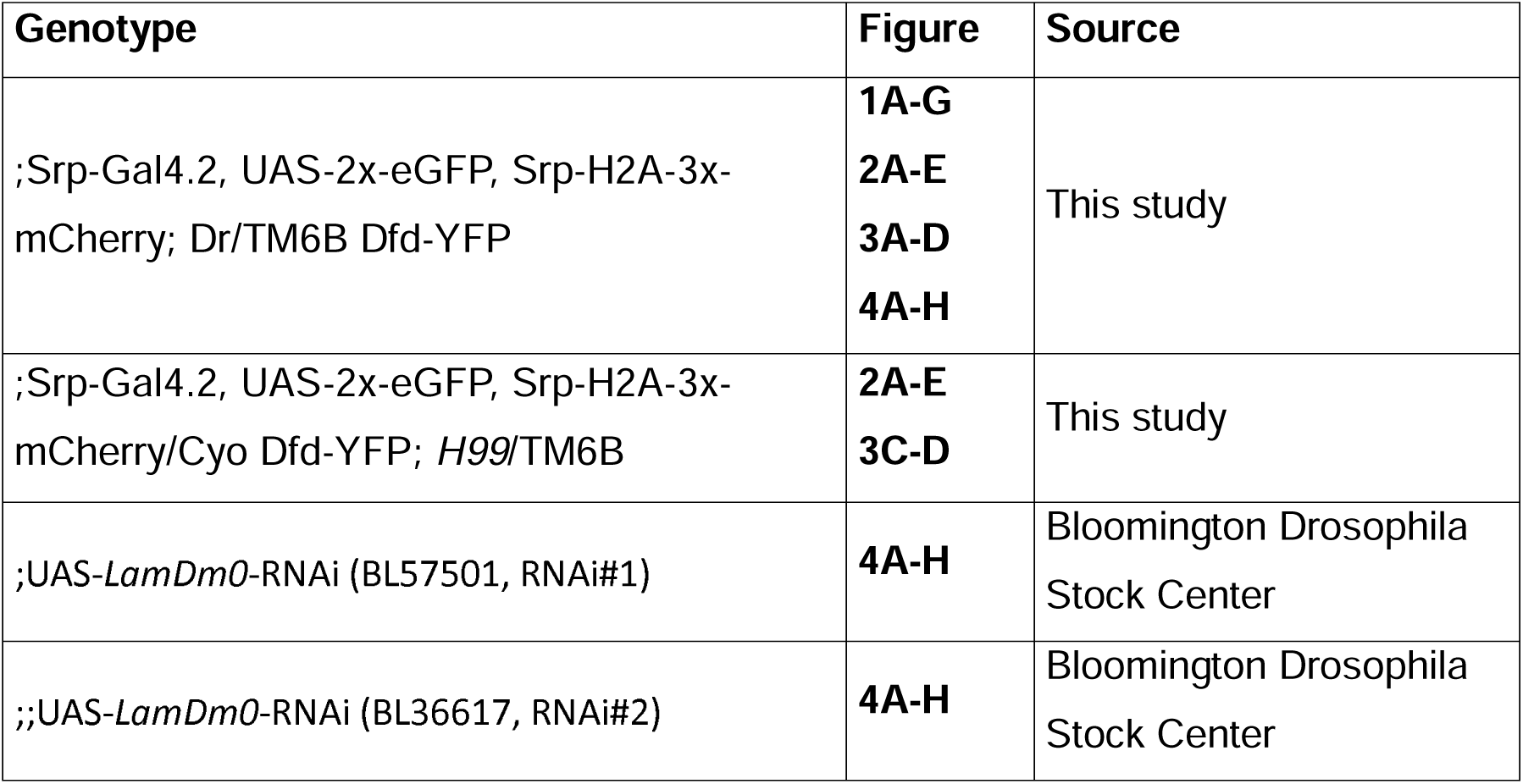
***Drosophila* strains**

**Table S2.** Statistics.

| Figure | N | Test | p-value |
| --- | --- | --- | --- |
| 1C | N<sub>Stage12</sub>: 5 embryos, 157 MΦs<br>N<sub>Stage15</sub>: 5 embryos, 64 MΦs | Unpaired t-test | p<sub>Stage12 vs Stage15</sub>= 0.0002 |
| 1E | N<sub>Stage12</sub>: 5 embryos, 69 MΦs<br>N<sub>Stage15</sub>: 5 embryos, 63 MΦs | Mann-Whitney test | p<sub>Stage12 vs Stage15</sub>= 0.0079 |
| 1F | N<sub>Stage12</sub>: 5 embryos, 81 MΦs, 113 indenting vacuoles<br>N<sub>Stage15</sub>: 5 embryos, 57 MΦs, 143 indenting vacuoles | Unpaired t-test | p<sub>Stage12 vs Stage15</sub>= 0.0054 |
| 1G | N<sub>Stage12</sub>: 5 embryos, 81 MΦs, 113 indenting vacuoles<br>N<sub>Stage15</sub>: 5 embryos, 57 MΦs, 143 indenting vacuoles | Unpaired t-test | p<sub>Stage12 vs Stage15</sub>= 0.1455 |
| 2D | N<sub>Control</sub>: 5 embryos, 63 MΦs<br>N<sub>H99</sub>: 5 embryos, 67 MΦs | Mann-Whitney test | p<sub>Control vs H99</sub>= 0.0079 |
| 2E | N<sub>Control</sub>: 5 embryos, 88 MΦs<br>N<sub>H99</sub>: 5 embryos, 43 MΦs | Unpaired t-test | p<sub>Control vs H99</sub>= 0.8401 |
| 3D | N<sub>Control</sub>: 5 embryos, 99 MΦs<br>N<sub>H99</sub>: 5 embryos, 100 MΦs | Unpaired t-test | p<sub>Control vs H99</sub>= 0.0020 |
| 4B | N<sub>Control</sub>: 5 embryos, 53 MΦs, 237 vacuoles<br>N<sub>LamDm0 RNAi#1</sub>: 10 embryos, 112 MΦs, 458 vacuoles<br>N<sub>LamDm0 RNAi#2</sub>: 10 embryos, 120 MΦs, 535 vacuoles | One-way ANOVA | p<sub>Control vs N<sub>LamDm0 RNAi#1</sub></sub>= 0.7049<br>p<sub>Control vs N<sub>LamDm0 RNAi#2</sub></sub>= 0.7573 |
| 4C | N<sub>Control</sub>: 5 embryos, 53 MΦs, 237 vacuoles<br>N<sub>LamDm0 RNAi#1</sub>: 10 embryos, 112 MΦs, 458 vacuoles<br>N<sub>LamDm0 RNAi#2</sub>: 10 embryos, 120 MΦs, 535 vacuoles | One-way ANOVA | p<sub>Control vs N<sub>LamDm0 RNAi#1</sub></sub>= 0.0470<br>p<sub>Control vs N<sub>LamDm0 RNAi#2</sub></sub>= 0.0012 |
| 4E | N<sub>Control</sub>: 5 embryos, 57 MΦs, 143 indenting vacuoles<br>N<sub>LamDm0 RNAi#1</sub>: 10 embryos, 119 MΦs, 227 indenting vacuoles<br>N<sub>LamDm0 RNAi#2</sub>: 10 embryos, 140 MΦs, 229 indenting vacuoles | One-way ANOVA | p<sub>Control vs N<sub>LamDm0 RNAi#1</sub></sub>= 0.0291<br>p<sub>Control vs N<sub>LamDm0 RNAi#2</sub></sub>= 0.0002 |
| 4G | N<sub>Control</sub>: 5 embryos, 63 MΦs<br>N<sub>LamDm0 RNAi#1</sub>: 10 embryos, 103 MΦs<br>N<sub>LamDm0 RNAi#2</sub>: 10 embryos, 128 MΦs | Kruskal-Wallis test | p<sub>Control vs N<sub>LamDm0 RNAi#1</sub></sub>> 0.9999<br>p<sub>Control vs N<sub>LamDm0 RNAi#2</sub></sub>= 0.0743 |
| **4H** | N<sub>Control</sub>: 5 embryos, 88 MΦs<br>N<sub>LamDm0 RNAi#1</sub>: 10 embryos, 93 MΦs<br>N<sub>LamDm0 RNAi#2</sub>: 10 embryos, 79 MΦs | One-way ANOVA | p<sub>Control vs N<sub>LamDm0 RNAi#1</sub></sub> = 0.0205<br>p<sub>Control vs N<sub>LamDm0 RNAi#2</sub></sub> = 0.0116 |
| **S1A** | N<sub>Stage12</sub>: 5 embryos, 157 MΦs<br>N<sub>Stage15</sub>: 5 embryos, 64 MΦs | 2way ANOVA | p<sub>Stage12:0 vs Stage15:0</sub> <0.0001<br>p<sub>Stage12:1-3 vs Stage15:1-3</sub> = 0.9728<br>p<sub>Stage12:4-6 vs Stage15:4-6</sub> = 0.0111<br>p<sub>Stage12:7-9 vs Stage15:7-9</sub> = 0.9946 |
| **S1B** | N<sub>Stage12</sub>: 5 embryos, 188 MΦs<br>N<sub>Stage15</sub>: 5 embryos, 88 MΦs | Unpaired t-test | p<sub>Stage12 vs Stage15</sub> = 0.0209 |

## Supplementary Movies

**Movie S1.**

3D rendering of a stage 12 (left) and a stage 15 (right) control embryos expressing a nuclear (Srp-H2A-3x-mCherry, magenta) and a cytosolic (Srp-Gal4.2, UAS-2x-eGFP, green) MΦ marker. Related to Figure 1A.

**Movie S2.**

3D rendering of a stage 12 (left) and a stage 15 (right) control MΦs expressing a nuclear (Srp-H2A-3x-mCherry, magenta) and a cytosolic (Srp-Gal4.2, UAS-2x-eGFP, green) marker. Related to Figure S1A.

**Movie S3.**

Time-lapse showing a stage 12 (left) and a stage 15 (right) control MΦs expressing a nuclear (Srp-H2A-3x-mCherry, magenta) and a cytosolic (Srp-Gal4.2, UAS-2x-eGFP, green) MΦ marker. Related to Figure 1D.

**Movie S4.**

Time-lapse showing a stage 15 control (left) and H99 mutant (right) MΦs expressing a nuclear (Srp-H2A-3x-mCherry, magenta) and a cytosolic (Srp-Gal4.2, UAS-2x-eGFP, green) MΦ marker. Related to Figure 2C.

**Movie S5.**

3D rendering of a stage 15 control embryo expressing a nuclear (Srp-H2A-3x-mCherry, magenta) and a cytosolic (Srp-Gal4.2, UAS-2x-eGFP, green) MΦs marker fixed and stained with an anti-Lamin Dm0 antibody (grey). Related to Figure 3A.

